# Endothelial mechanobiology under controlled disturbed flows

**DOI:** 10.1101/2023.08.27.555040

**Authors:** Neha Paddillaya, Suyog Mahulkar, Jaywant H Arakeri, Namrata Gundiah

## Abstract

Early atherosclerotic lesions often develop in areas of disturbed flow within arterial curvatures and bifurcations. Complex flows alter the signaling of inflammatory and other signaling molecules that may exacerbate the disease phenotype. Microfluidic platforms to assess changes in endothelial mechanobiology generally employ laminar unidirectional or oscillatory flows generated in straight channels; such platforms do not however mimic the complex time-varying bi-directional shear patterns on arterial walls. We fabricated an endothelium-on-chip device to generate “controlled” disturbed flows, characterized using a wall shear rosette, such as those reported in aneurysmal vessels and in regions with atherosclerotic plaques. We cultured human aortic endothelial cells (HAEC) in the device and subjected the monolayer to a circular shear rosette which represents bidirectional and oscillatory shear stresses. Immunofluorescence results show large cuboidal cells with significantly higher nuclear areas, changes in the localization of VE-Cadherin, elevated actin and NF-kB expressions in the device as compared to cell monolayers subjected to laminar flow, unidirectional oscillatory flow, and No-flow conditions. The creation of a dysfunctional endothelial monolayer due to bidirectional oscillatory flows also correlated with dramatic changes to the lamin A/C distribution and heterochromatin organization in the nucleus that have not been reported earlier. Such studies are potentially useful to assess novel therapeutics, mitigate effects of vascular disease for personalized medicine, study thrombus creation in the vicinity of inflamed endothelial monolayers, and reduce ourreliance on animal trials. Our device and analytical method represent the first successful demonstration of generating controlled disturbed flows in microfluidic devices

## 1.0 Introduction

Endothelial cells (EC) line the blood vessel lumen and experience cyclically varying complex shear stresses due to the blood flow. Wall shear stress (WSS) activates multiple signal transduction pathways through adhesive proteins that link the EC monolayer to the underlying matrix, cytoskeletal proteins, caveolae, tyrosine kinase receptors, ion channels, the glycocalyx, and primary cilia.^1^ Response of the endothelium to mechanical stimuli is important in angiogenesis, vascular remodeling, and chronic inflammatory arterial diseases, including atherosclerosis.^2–7^ Straight sections of small arteries and capillaries generally experience laminar shear stresses, whereas branch points, regions downstream of curved sections, and bifurcations are subjected to more complex stresses during normal cardiac activity **(Figure 1a and 1b)**. Early atherosclerotic lesions develop along curved regions and in arterial bifurcations.^8,9^ Macrophages localize below the endothelium in regions of lower curvature in the murine aortic arch and at branch points of large arteries.^10^ Higher atherosclerotic expression in regions experiencing disturbed flows correlates with changes in WSS magnitude and direction.^11,12^ Complex flows activate and upregulate endothelial NO synthase (eNOS) *via* inflammatory signaling due to NF-kB.^10–13^ Increased nitrous oxide (NO) results in higher expression of leukocyte adhesion molecules, including VCAM-1 and E-selectin, that promote the diseased phenotype.^10^

**Figure 1:**
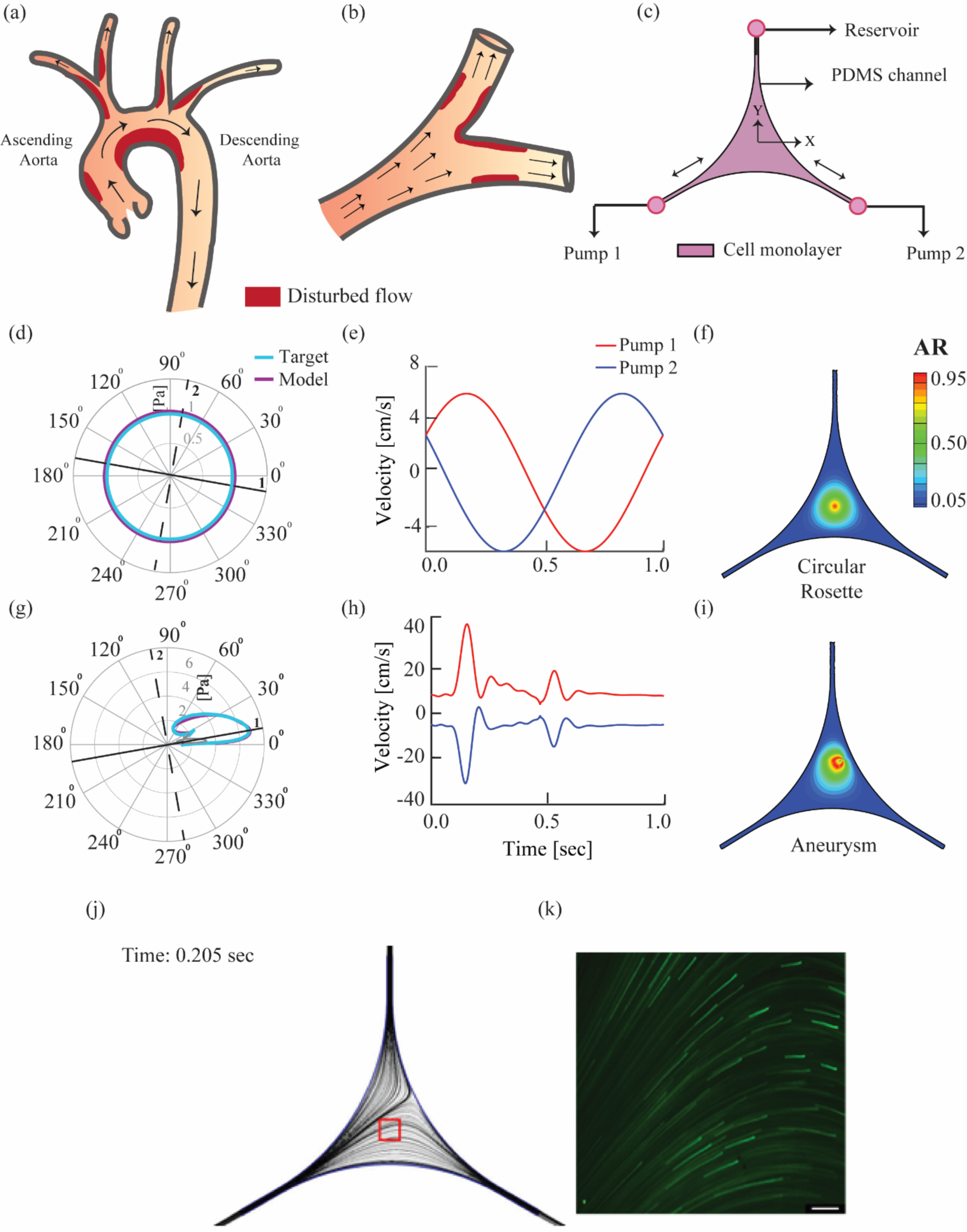
Disturbed flow regions are shown (red) in the **(a)** aortic arch and descending artery, and **(b)** arterial bifurcations that experience secondary flows which contribute to the build-up of atherosclerotic plaques. **(c)** A cartoon of the novel endothelium-on-chip device, fabricated using PDMS, is shown that includes the HAEC monolayer in the device. Inlet channels to the device were connected to two syringe pumps. **(d)** Rosettes are polar plots that show variations in shear stresses at each location in the device over one flow cycle. The target circular rosette profile (cyan) closely matches the rosette obtained computationally using velocity inputs (dark blue) from solutions to the semi-analytical model. **(e)** Inlet velocity profiles for the two pumps were used to generate a circular shear rosette in the device. **(f)** The contour plot shows variations in anisotropy ratio (AR), calculated as a ratio of the minor to major axes lengths, for a circular rosette (AR=1) at the device centroid. **(g)** Target shear rosette (cyan) for a point in the inner carotid artery (ICA), with aneurysm^15^, is shown along with the rosette calculated usingthe model (dark blue). **(h)** Corresponding velocities to the pump inlets from the semi-analytical model, and **(i)** AR contour map at the device centroid are shown for the ICA aneurysm. **(j)** Transient streamlines from CFD calculations are shown at a time point of 205 ms **(k)** Corresponding streak images, indicating streamlines from flow visualization experiments, are shown for comparison. Scale bar 100µm.

Physiological arterial flows are significantly more complex than laminar and oscillatory flows due to the unsteady or pulsatile and spatially developing conditions.^14–16^ High wall curvatures, such as those in the ascending aorta and smaller cerebral arteries, lead to the generation of secondary flows which are caused by an imbalance between centrifugal forces and the radial pressure gradient.^16^ Interactions between unsteadiness and curvature are highly nonlinear and result inflow separations and reversals, vortical structures, and regions of WSS stagnation/ fixed points that are broadly labeled as “disturbed” flows.^16^

Devices based on microscale technologies mainly use laminar unidirectional or oscillatory flows in straight microfluidic channels to assess changes in endothelial mechanobiology.^11,17–21^ Generating controlled disturbed flows, similar to those found in disease prone arterial regions, is an ongoing challenge to implement in lab-on-chip devices.^22^ Whereas steady laminar channel flow past a ridge/ obstruction produces a separated flow in the wake, no secondary flows are generated.^21,23,24^ Alternate methods, such as orbital shaker, do generate complex multidirectional shear stresses that can be applied to EC monolayers.^25^ Such methods are, however, not capable of creating specified and controlled temporal variations in WSS.

We recently proposed a graphical method to characterize the temporal changes in flow dynamics at each point of the arterial wall. This method uses variations in the WSS vector along the principal and orthogonal flow directions, and represents them as a polar “shear rosette” plot.^15^ The rosette size is a measure of the WSS magnitude, variations about the centroid indicate oscillations, and shape deviations from linearity signify the extent of flow bidirectionality.^15^ The ratio of the rosette width to length, defined as anisotropy ratio (AR), is a useful method to quantify secondary flows. For example, a circular rosette has AR=1 represents a case where the WSS magnitude is constant but its direction changes through 360 degrees in one cycle. Regions with high AR values, such as the inner curvature of ascending aorta and arterial bifurcations, exhibit strong secondary flows, whereas those with AR=0, such as straight sections of arteries, experience oscillatory, unidirectional flows. The AR metric is hence useful to quantify the presence and strength of secondary flows in arteries.

In this study, we fabricated a microfluidic device, employing two synchronized syringe pumps, to create customized shear rosette patterns at the device centroid. To achieve this customization, we developed a semi-analytical model to calculate variations in time of the input flow rates to each pump and validated the model accuracy for various rosette shapes using computational fluid dynamics (CFD) and experimental flow visualization.We studied changes in cellular and nuclear mechanobiology using human aortic endothelial cell (HAEC) monolayers subjected to a circular shear rosette in the device. We compared these results with HAEC monolayers exposed to laminar and oscillatory unidirectional flows in straight microfluidic channels; cells on coverslips that were not under flow were used as controls.PCR studies show several fold changes in inflammatory NF-kB, VE-Cadherin, and lamin A/C markers. Heterochromatin condensation was clearly visible in cells subjected to oscillatory flows. Elucidating the mechanisms underlying EC responses to controlled disturbed flows is essential to our understanding of factors that may be involved in the pathophysiology of vascular disease, developing patient specific therapeutics to modulate EC behaviors, assessing bacterial/ viral interactions with EC, and characterizing thrombus formation.

## 2.0 Results and Discussion

### 2.1 Generation of “controlled” disturbed flows using microfluidics

We fabricated an endothelium-on-chip device using replica molding with PDMS **(Figure 1c).** The device consists of three arms with an angular separation of 120° **(Figure 1c).** Two of these arms are connected to synchronized syringe pumps that serve as inlet ports to the device, whereas the third arm is a reservoir port. We adjusted the input flow rates to the two pumps using the semi-analytical model (see Methods) to achieve any desired shear rosette pattern at the device centroid. For a physiological WSS magnitude of 1Pa^24^, the inlet flow velocities required to generate a circular rosette at the device centroid **(Figure 1d)** are given by

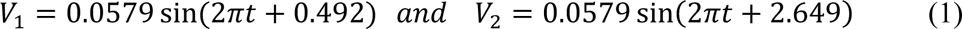

These expressions are different when used to generate other rosette shapes. We validated this approach with CFD implemented using ANSYS-Fluent **(Figure S1 and S2, Table S1 and S2).**

Figure 1d shows a significant overlap between rosette shapes from CFD simulations with the targeted circular rosette at the device centroid. Velocities from Equation 1 were used at the pump inlets, and the corresponding WSS variations were calculated (Figure 1e**; Movie S1).** We computed the AR values at each point in the device using temporal variations in WSS. Figure 1f shows a contour plot with radially varying AR that changes in magnitude with distance from the device centroid. Regions close to the inlets and the reservoir have WSS regions characterized by AR=0 in the device.

The semi-analytical model is useful to generate any arbitrary shear rosette shape to describe disturbed flow conditions at different regions in the circulatory system. Figure 1g shows replication of the rosette for diseased aneurysmal inner carotid artery (ICA) with model calculations at the device centroid that were obtained using inlet velocities in Figure 1h.^15^ Peak AR values (=0.45), shown as a contour map in Figure 1i, are off-centered from the centroid for this condition. Movie S2 shows time variations in the inlet pumps that produce the targeted rosette for ICA. We replicate a few other conditions, such as an elliptical **(Figure S3a)** and a complex rosette **(Figure S3d)**, reported in an aneurysmal ascending aorta, using this method.^15^ To experimentally confirm these results, we performed flow visualization using fluorescent microspheres as tracer particles for a circular rosette condition **(Movie S3).** The tracer particle streak images, obtained with long camera exposure, show flow streamlines within the channel. A comparison between streamlines obtained from CFD studies and streak images fromflow visualization experiments is shown in **Figures1j and 1k (Movie S3).** These streamlines show a good match with those obtained using CFD calculations. The model hence presents a robust approach to generate arbitrary shear rosette conditions that characterize disturbed flows.

Various methods have been used to simulate disturbed flow conditions in diseased arterial regions. These include naturally occurring or surgical interventions in animal models, vertical step flow chambers, modified cone-plate viscometers, and microfluidic systems to elucidate the phenotypic adaptation of cells to flow.^17–20^ Animal models for atherosclerosis often involve partial ligation of arteries that alter flows and cause changes incellular behaviors from atheroprotective to atheroprone in ligated regions.^26^ Developing therapeutics to manage and mitigate vascular disease is a continuing exercise due to the high costs for animal studies in screening novel compounds. *In vitro* experiments suggest inclusion of substrate stretching with unidirectional laminar flows to better represent the *in vivo* cellular milieu.^27,28^ Assays that preserve the physiological complexity are clearly warranted but are challenging to implement due to limited access to imaging at cellular and sub-cellular levels. There is a need for lab-on-chip platforms that accurately capture the complex biomechanical milieuto test a vast repertoire of flow conditions, test cellular responses to specific treatments, and to better understand the development of vascular diseases. Our device and analytical method represent the first successful demonstration of generating controlled disturbed flows in microfluidic devices.^29^

### 2.2 Endothelial cells are enlarged and rounded under bidirectional oscillatory flows

EC morphologies are spindle shaped and align in the flow direction under laminar and unidirectional conditions in contrast to atheroprone regions that have rounded cells.^5,7,30–32^ How do bidirectional oscillatory flows influence endothelial cellular and nuclear morphologies, is an important question. To explore possible changes in EC morphologies, we cultured the HAEC monolayer on a collagen-1 coated coverslip and clamped it to the PDMS device (see Methods). This method significantly reduces the time required to grow EC monolayers in microfluidic devices and minimizes the media used to achieve cell confluency prior to performing the flow studies. Finally, we added inlets to the pumps and a reservoir to create the endothelium-on-chip device.^29^

We subjected the HAEC monolayer to a circular shear rosette (Bi-O) in the device for 24 hours and stained the monolayer with Rhodamine phalloidin to visualize actin (red) and DAPI to label the nuclei (blue). We used the TissueAnalyser plugin in Fiji (ImageJ) to segment cell boundaries and nuclei in the monolayer, and calculated the correspondingareas and aspect ratios (∼300 cells) (Figures 2a**, 2b, and 2c).** We compared results from this flow condition with HAEC monolayers in straight channels subjected to unidirectional laminar (Uni-L) and oscillatory (Uni-O) flows, and used the monolayer under No-flow condition as a control in this study (**Table 1**). Our results show that laminar flows resulted in highly elongated morphologies along the flow direction. HAECs in the Uni-O group had the lowest cell areas and aspect ratios as compared to the Uni-L group that had the corresponding highest values (Figure 2d). Areas for the No-flow group were lower than Uni-L and higher than Uni-O groups, whereas the aspect ratios for this group were not significantly different from the Uni-O group (Figure 2d **and 2e).** The Bi-O group had cuboidal cells with significantly greater areas as compared to the Uni-O and No-flow groups (p<0.05). These results demonstrate that HAEC monolayers remodel through changes in their cytoskeletal and nuclear morphologies in response to fluid shear stress.

**Figure 2:**
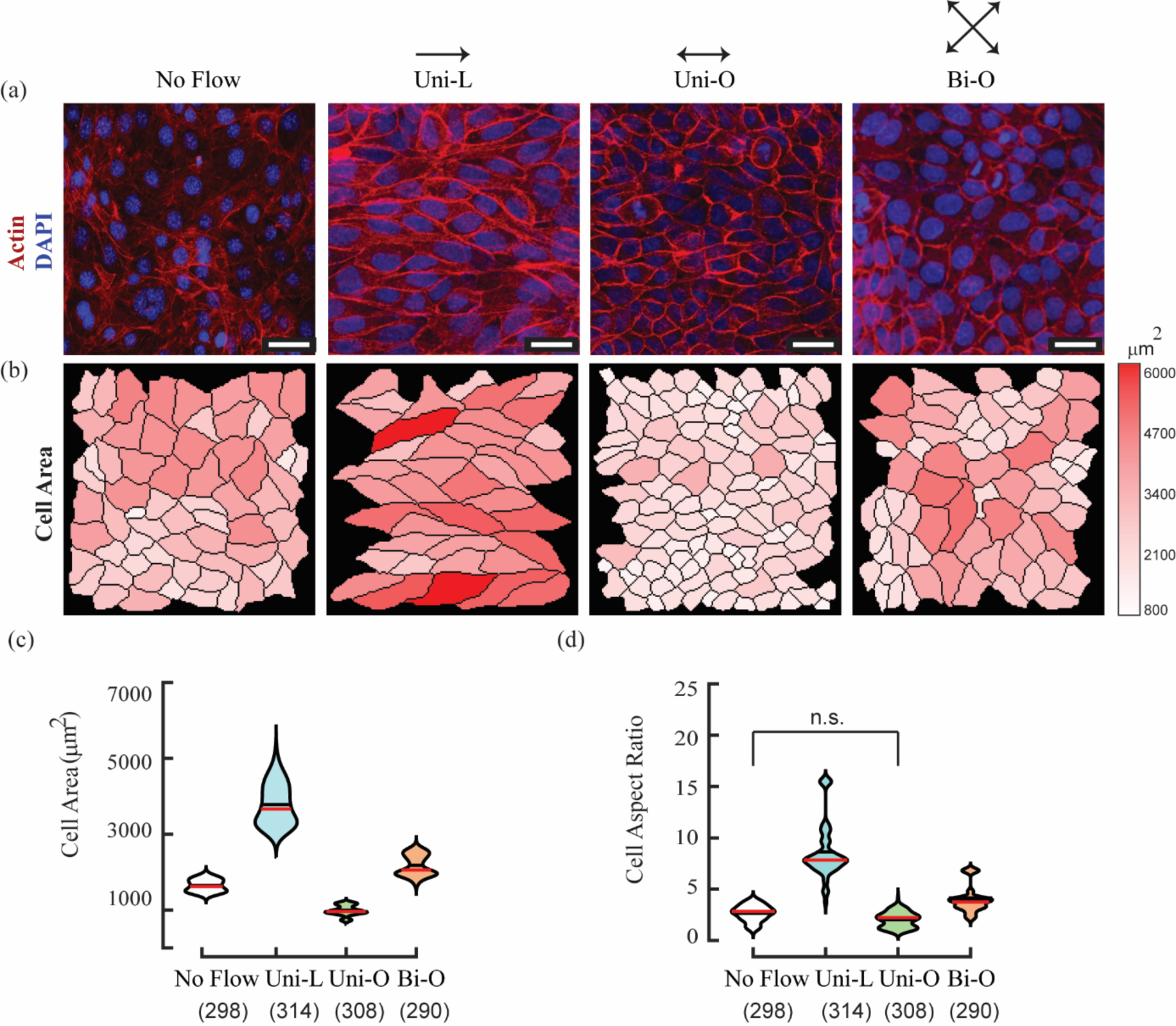
HAEC remodel differentially underflow (a) Cell morphologies in response to No-flow (Control), Uni-L, Uni-O, and Bi-O conditions subjected for 24 hours are shown. Actin was stained with Rhodamine phalloidin (red) and the nucleus with DAPI (blue). Scale bar 25µm. (b) Cell boundaries and (c) Cell areas were obtained using TissueAnalyser (ImageJ). Violin plots show differences in (d) cell areas and (e) aspect ratios for the different flow conditions. Number of cells in each group (N) is indicated within parenthesis in the figures. The means for each group are shown in red. ANOVA with Bonferroni tests showed significant differences (p < 0.05) in group comparisons with the exception of those marked “n.s.” in the plots.

**Table 1:** Morphological differences in HAEC monolayers for the different flow conditions in this study.

|  | *No-flow* | *Uni-L* | *Uni-O* | *Bi-O* |
| --- | --- | --- | --- | --- |
| ***Cell area [$\mu\text{m}^2$]*** | 1642.81<br>$\pm 155.23$ | 3491.61<br>$\pm 655.90$ | 996.93<br>$\pm 131.78$ | 2177.12<br>$\pm 270.34$ |
| ***Aspect Ratio*** | 2.67<br>$\pm 0.75$ | 8.63<br>$\pm 2.64$ | 2.01<br>$\pm 0.73$ | 3.98<br>$\pm 1.27$ |
| ***Nuclear area [$\mu\text{m}^2$]*** | 230.23<br>$\pm 23.95$ | 318.45<br>$\pm 28.64$ | 287.79<br>$\pm 29.94$ | 387.74<br>$\pm 74.33$ |
| ***Cell: Nucleus areal fraction*** | 7.22<br>$\pm 1.06$ | 11.12<br>$\pm 2.50$ | 3.50<br>$\pm 0.60$ | 5.83<br>$\pm 1.38$ |
| ***Actin Intensity [AU]*** | 18.19<br>$\pm 1.39$ | 32.72<br>$\pm 3.03$ | 21.51<br>$\pm 1.83$ | 18.60<br>$\pm 3.35$ |
| ***VE-Cadherin Intensity [AU]*** | 17.99<br>$\pm 2.41$ | 32.72<br>$\pm 3.03$ | 15.46<br>$\pm 0.97$ | 22.19<br>$\pm 1.20$ |

ECs modify their morphology and orientation in response to flows.^30^ Cellular morphologies *in vivo* are dramatically different at branch points of mouse and primate aorta that experience disturbed blood flows as compared to laminar flow regions in straight arteries.^24,33^ Cells in branch points are cuboidal as compared to elongated morphologies in the straight regions of the arteries.^24,33^ Results from our study also confirm that cells under laminar flows had elongated morphologies and high actin intensity in contrast to those which were subjected to oscillatory flows that had cuboidal cells.

The seeding density was maintained the same at the start of all experiments in our study. Interestingly, results from our study show clear differences in cell sizes for all groups at 24 hours after the application of flow. Chien and coworkers examined the luminal surface of the rabbit thoracic aorta in disturbed flow regions and showed that EC have higher mitotic rates.^34^ Disturbed flows cause an increase in the endothelial cell proliferation through the downregulation of p53 and inhibition of AKT in mouse and pig aorta.^32^ Increased cell senescence *via* p53- and sirtuin-1-dependent pathway have also been reported.^32,34^ In contrast, laminar flows are known to suppress the cell cycle progression by upregulation of cyclin-dependent kinase inhibitor 1 (p21).^35^ These studies hence demonstrate G0/G1 phase arrest under laminar flow conditions whereas disturbed flows promoted cell cycle progression (S+G2/M phase).^35,36^ The differences in cell sizes in the Uni-O and Bi-O groups in our study may be a result of cell cycle arrest or increased proliferation. Additional studies are however warranted to delineate possible differences in proliferations in EC monolayers subjected to bidirectional oscillatory flows.

### 2.3 VE-Cadherin disruptions are augmented in monolayers under bidirectional oscillatory flows

Endothelial barrier function is maintained by junctional proteins that regulate the transport of biomolecules.^37^ Membrane permeability is critical for monolayers to preventbacterial/ viral infections, inflammation, cancers, diabetes, atherosclerosis, and other diseases.^38,39^ VE-Cadherin mediated contacts and VEGF-R2 complexes, including PECAM, are known to be critical transducers of shear stresses exerted on EC monolayers.^37,39,40^ Based on the dramatic differences in the cell morphologies under various flow conditions in our study, we asked if there were any changes in the junctional proteins of HAEC monolayers in the different flow groups. We used immunofluorescence methods to investigate changes in VE-Cadherin and stress fiber (SF) formation under shear (see Methods).

Representative images (Figure 3a) from the groups show clear differences in the presence and distribution of VE-Cadherin in monolayers subjected to laminar flow (Uni-L) as compared to other groups. VE-Cadherin stained continuously at the cell borders in the Uni-L group, whereas the Bi-O group showed discontinuous expression.

**Figure 3:**
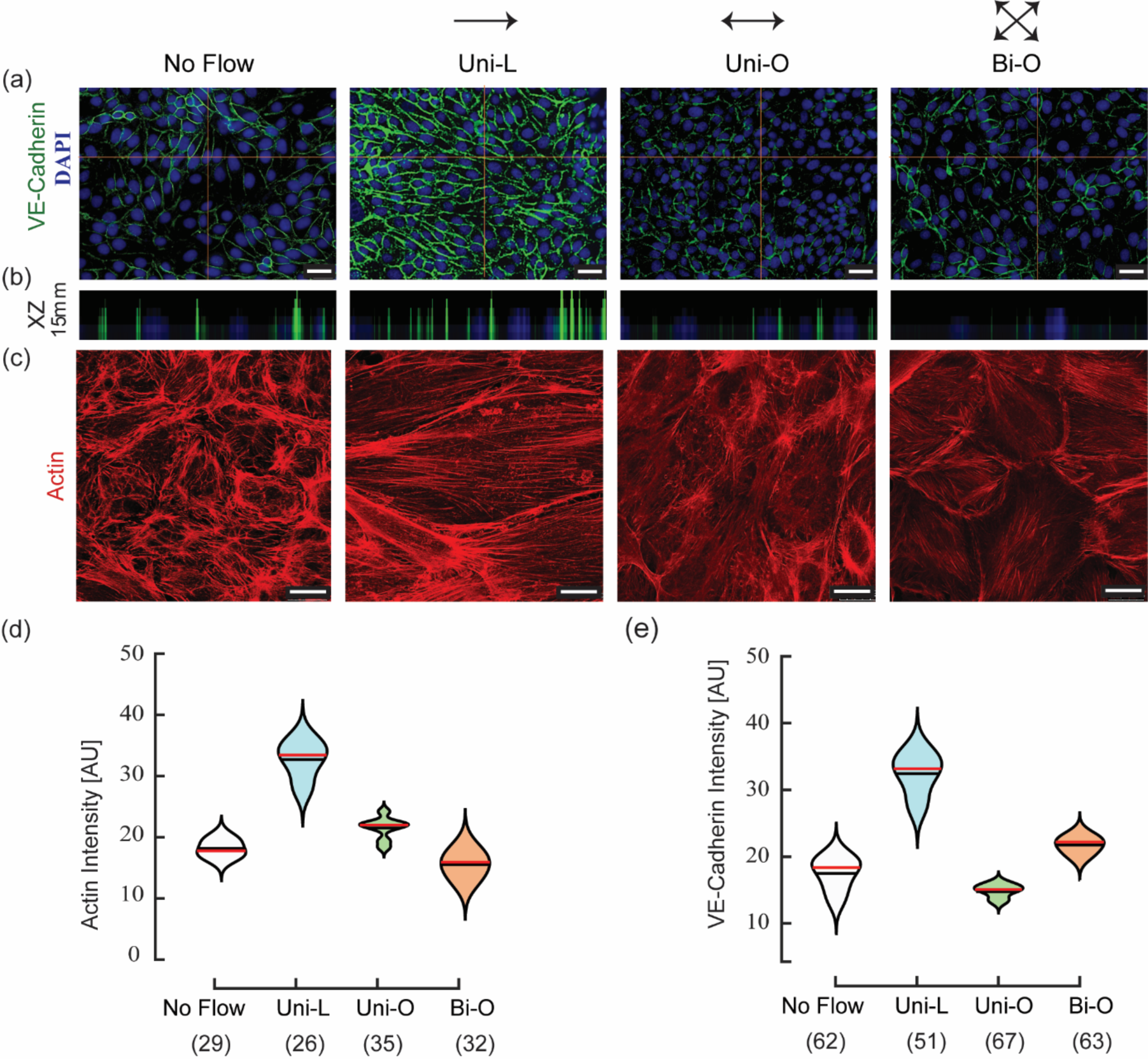
Changes in VE-Cadherin and actin are visible due to differential flow conditions **(a)** Immunofluorescence images of HAEC monolayers are shown for VE-Cadherin (green) and nucleus (blue). Scale bar 25 µm. **(b)** XZ image for the VE-Cadherin image at the cross-hair point labeled in **(a)** shows uniform expression of cell junctions for the Uni-L group as compared to the other groups. **(c)** Immunofluorescence images for actin (Red) show differences in stress fibers for the various groups. Scale bar 25 µm. These images were used to quantify the intensity of **(d)** Actin and **(e)** VE-Cadherin for the various groups. Number of cells in each group is indicated within parenthesis in the figure. ANOVA with Bonferroni tests showed significant differences (p < 0.05) for all groups.

We also see reduced fluorescence intensity in the oscillatory (Uni-O and Bi-O) and control (No-flow) groups, respectively. XZ sectional images for VE-Cadherin (Figure 3b) demonstrate differences in monolayer heights under oscillatory flows, No-flow, and Uni-L conditions. Figure 3c shows well-defined SFs that were aligned along the flow direction under unidirectional laminar conditions as compared to cuboidal cells in the Bi-O group which had few aligned SFs along the flow direction. We used these images to quantify actin intensities in monolayers subjected to various flow conditions (**Table 1**; Figure 3d). There were significant differences in actinintensities for the Bi-O group as compared to the Uni-O group (p<0.05). VE-Cadherin expression was higher in the Uni-L group as compared to all other groups (Figure 3e). In contrast, VE-Cadherin was lowest for the Uni-O group and was partially lower in the Bi-O group. There were no significant differences between the No-flow and Uni-O groups (Figure 3e).

Earlier studies have demonstrated reduced monolayer permeability under laminar shear stress through occludin and adherens junctions.^41^ Results from our study show that unidirectional oscillatory flows significantly lowered VE-Cadherin, whereas bidirectional oscillatory flows led to partial downregulation of VE-cadherin. A reduction in membrane-bound VE-Cadherinin disturbed flow regions leads to vascular destabilization, intercellular gap opening, loss of barrier function, and a correspondingly enhanced macromolecular permeability.^37^ Disruption and reduction in VE-Cadherin expression under oscillatory conditions hence suggesta possible redistribution of gap junctions and cytoskeletal proteins that can compromise the monolayer integrity and increase the membrane permeability.^42^ Our data suggest that redistribution of cytoskeletal proteins due to bidirectional oscillatory flows mayalterthe membrane permeability and tractional stresses,controlled by ventral stress fibers, to change cell-substrate adhesions. A measurement of the monolayer permeability alone may not represent changes in the mechanobiological behaviors of endothelial cell monolayers. An understanding of the causal links between morphology, VE-Cadherin, SF formation, and inflammation in endothelial monolayers can be tested in future studies using *in vitro* microfluidic platforms.

### 2.4 Nuclear membrane disruptions and heterochromatin condensation characterize monolayers subjected to oscillatory flows

Mechanical forces regulate the nuclear morphology, organization of chromatin, and gene regulation.^43–45^ The application of shear stress elicits several notable effects, including anincrease in stiffness, reduction in cellular height, alignment of cellular nuclei along the axis of flow, and remodeling.^46^ Laminar flows are accompanied with nuclear alignment and remodelingthat are hypothesized to minimize the effects of the shear stress.^47^ Heterochromatin condensation spotswithin the nucleus in response to mechanical stress indicate changes in the chromatin organization of cells.^48,49^ Because cellular responses are critical in mitigating theirexposure to mechanical forces and preserving genomic integrity, we asked if there were any visible changes in the cell nuclei when exposed to Bi-O flows. DAPI stained nuclear images (Figure 4a) in our study demonstrate distinct differences in the organization of heterochromatin structures, visible as multiple bright spots, in Uni-O and Bi-O monolayers compared to amore homogenous nuclear staining in the other groups.

**Figure 4:**
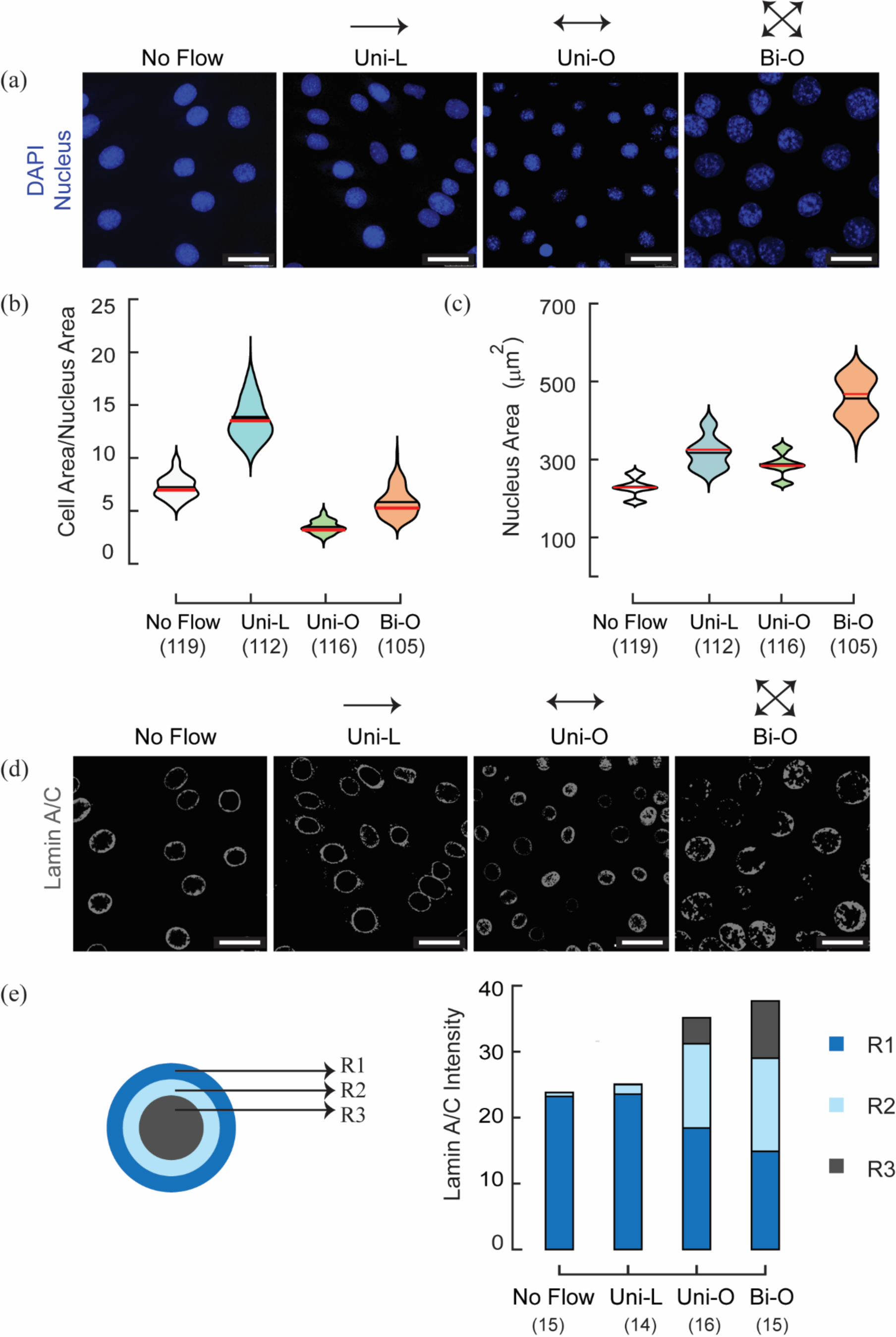
Chromatin condensation and Lamin A/C disruption **(a)** Immunofluorescence images of DAPI stained (blue) nuclei in HAEC monolayers are shown for the various groups in our study. Heterochromatin condensation is seen under oscillatory flows (Uni-O and Bi-O). **(b)** Nuclear areas were obtained for the various groups in our study and **(c)** Cell area to nuclear area ratio. **(d)** Immunofluorescence images show disruption in the localization of Lamin A/C in the nuclei under oscillatory flow conditions (Uni-O and Bi-O). Scale bar is 25µm. ANOVA and Bonferroni test demonstrated significant differences (p < 0.05) between all group comparisons.**(e)** We used a graphical representation to quantify the differences in Lamin A/C redistribution within three regions of equal area in the nucleus. Cells under oscillatory flows (Uni-O and Bi-O) have lamin distributed away from the edge of the nuclear membrane. Number of cells in each group are included within the parenthesis in these plots.

Results show that cells in nuclear areas in Bi-O cells were 68.41% larger (p<0.05) compared to those in the No-flow control group (Figure 4b). The Uni-L group had significantly higher cytoskeletal/ nuclear area ratios, followed by No-flow, Bi-O, and Uni-O groups (Figure 4c; **Table 1).** The role of the nuclear lamina in endothelial mechanotransduction under differential shear stresses is poorly understood. Lamin A/C in the nuclear lamina is reported to act as a scaffold for heterochromatin and other transcriptional proteins, and is critical in preventing force induced DNA damage^48,50–52^, we hence assessed for possible changes in the localization of lamin A/C in our experiments that maybe altered under oscillatory flow conditions (Figure 4d).

Endothelial monolayers showed distinct disruption and mislocalization of LaminA/C in the nuclear mid-section, occurring from the membrane to the interior, in HAEC monolayers subjected to oscillatory flows. In contrast, cells in No-flow and Uni-L groups had Lamin A/C localization at the nuclear envelope.The nuclear lamina not only maintains the nuclear shape but also plays a critical role in the bidirectional transmission of forces between the cytoskeleton and the nucleus.^53^

To quantify the Lamin A/C disruption in EC monolayers, we divided the nuclear boundary into three concentric regions (R1, R2, and R3) of equal areas from the edge (Figure 4e).Comparisons between the intensities of Lamin A/C in these regions show localization at the nuclear membrane (R1 region) for No-flow and Uni-L conditions.Lamin A/C intensity was however 16.8 times higher in the R2 and R3 regions for cells in the Bi-O group, and 10.9 times higher for the Uni-O group as compared to Uni-L. Loss of Lamin A/C under oscillatory flows correlated with the chromatin condensation in DAPI stained images of the endothelial cells (Figure 4a). These results suggest that disruption and mislocalization of Lamin A/C may potentially impact endothelial function at disturbed flow regions.

### 2.5 Inflammation and changes in gene expression levels under oscillatory flows

The pro-inflammatory transcriptional factor, NFκB, is implicated in many vascular diseasesincluding atherosclerosis and atherothrombosis.^54,55,54,55^ Activation of NF-κB triggers other inflammatory mediators that modulate smooth muscle cell activation; these may further contribute to inflammation and progression of the atherosclerosis phenotype.^33,54,56,57^ Studies demonstrate increased inflammation in ECs monolayers subjected to fluid flows acting perpendicular to aligned cellsas compared to those along the aligned monolayer direction that are anti-inflammatory and contribute to atheroprotection.^58^ Inflammatory markers are shown to be associated with the phosphorylation of Lamin A/C and a loss of the nuclear lamina.^52^ *In vitro* microfluidic platforms to create custom flow conditions can be useful to delineate these roles in endothelial monolayers. We next asked if there were changes in the gene expression levels in the monolayers subjected to the different flow conditions. We used PCR to analyze the regulation of mRNA levels for Actin, VE-Cadherin, NFκB, and LMNA genes in the various groups in our study and compared results with corresponding values in the No-flow group (Figure 5). Our results show increased NFκB and Lamin A/C expression under oscillatory flows. The 2.29±0.81 fold upregulation in LMNA gene expression in Bi-O specimens in our study may hence be a compensatory mechanism for reduced Lamin A/C at the nuclear lamina. Bi-O cells had 3.76±0.42 fold upregulation in the transcriptional factor, NFκB, as compared to other groups. NFκB expression was downregulated in the Uni-Lgroup (0.62±0.44), which clearly demonstrates the role of laminar unidirectional flows in anti-inflammatory response. LMNA gene expression was also upregulated in the Bi-O (2.29±0.81) group. There were higher fold changes in the expressions of VE-Cadherin and actin for the Uni-L group relative to Bi-O and Uni-O conditions. Laminar flows hence enhanced the VE-Cadherin gene expression in HAEC monolayers that broadly agree with the immunofluorescence data in our study (Figure 4b). The increased expression of inflammatory genes in the Bi-O groupin our study also agrees with results reported from *in vivo* studies of disturbed flow regions in mouse.^55^

**Figure 5:**
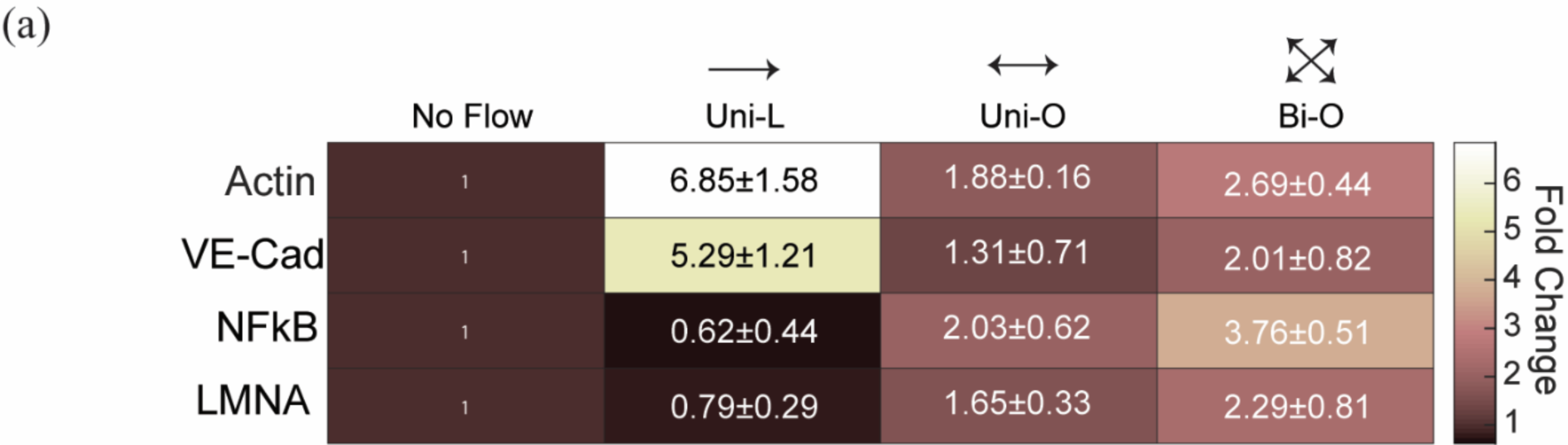
PCR quantification of mRNA fold changes are shown for Actin, VE-Cadherin, NFκB, and LMNA genes for the various groups in our study. Fold change in each group was normalized to GAPDH and No-flow conditions. Cells in the Bi-O group show elevated expression of all markers as compared to other groups. Data represents ± standard deviation from triplicates in the study.

Complex fluid flows, commonly occurring at branches or curves in the arterial system, promote misalignment and increased proliferation of endothelial cells with persistent low-grade inflammation.^59^ Such changes activate transcription factor complexes, such as NF-κB, AP-1, and YAP/TAZ, which lead to the upregulation of pro-inflammatory gene expression and increased cellular proliferation.^60,61^ There is hence a compelling rationale to investigate how endothelial nuclei perceive and respond to controlled disturbed flows, and to link these responses to changes in cell junction integrity and monolayer permeability. Changes to Lamins A/C, heterochromatin condensation, and increased inflammation in response to secondary or disturbed flows have not been reported earlier in endothelial studies to the best of our knowledge. These observations underscore the critical interplay between the nucleus and the cytoskeleton in vascular mechanics.

## 3.0 Conclusions

Mechanical forces are critical regulators ofthe vascular biochemistry, gene expression, and homeostasis in endothelial monolayers. Our work has four main implications. First, we developed a robust method to generate bidirectional oscillatory flows in a microfluidic device to replicate shear rosettes that characterize disturbed flows. The semi-analytical method computes inlet velocities to pumps in the device and is useful to generate any rosette shape at the device centroid. We demonstrate a good match between CFD calculations and flow visualization studies to validate our method. This approach can be used to generate complex rosettes reported in disease prone arterial regions.^15^ Second, we subjected HAEC monolayers to a circular shear rosette of 1Pa WSS magnitude, and compared morphological cellular changes with monolayers subjected to laminar unidirectional and oscillatory flows. Immunofluorescence studies show distinct differences in EC morphologies and actin intensities due to flows. Third, we observed disruption in Lamin A/C and reorganization of heterochromatin in the nuclei of endothelial cells in the Bi-O group; these changes were not visible in Uni-L and No-flow conditions. Finally, we measured changes in the gene expression levels and demonstrated significantly elevated actin, VE-Cadherin, NF-kB, and lamin expression in EC monolayers in the Bi-O group that have not been reported earlier. Together, these results demonstrate the ability of the endothelium-on-chip device to assess the effects of disturbed flows on endothelial mechanobiology. These studies also provide the potential to evaluate mechanosensory pathways implicated in conditions such as diabetes, and test therapeutics that target the vascular function. The device in this study represents a significant advance and provides a simple, versatile, inexpensive, and robust method to generate any disturbed flow conditions. The semi-analytical model may be used to generate any arbitrary rosette, to potentially simulate flow patterns observed in diseased arteries. Such devices can provide accurate, efficient, and reliable results to decrease the exorbitant costs of animal trialsand provide solutions for personalized medicine for a variety of genetic conditions. An understanding of how disturbed flows cause differential inflammation in endothelial mechanobiology, and the possible links to nuclear lamina disruption, may be useful in developing strategies to prevent and treat vascular dysfunction. Quantification of the extent of disruption of lamin expression may be useful in providing insights into the morphological and physiological implications of cellular responses to complex flows.

## 4.0 Methods

### 4.1 Development of a semi-analytical model to calculate flow velocitiesas inputs in the endothelium-on-chip device

The microfluidic device consists of three arms with an angular separation of 120° (Figure 1c). The device can be used to produce any shape of the WSS rosette, representing a polar plot of the temporal variations in WSS vectors, at the device centroid.^29^ This rosette may be regular, such as circular or elliptical shapes, or have complex shapes reported in diseased vessels.^15^ We use a laminar and quasi-steady assumption to compute the flow rates into the device inlets using a semi-analytical model. In this method, we use the centroid of the device as the origin of the coordinate system (Figure 1c). Based on the flow rates to the two pumps, we obtain the flow velocities at each of the two inlets, labeled *V*_1_and *V*_2_, as

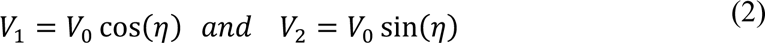

*V_0_* is the amplitude, and *η* is the phase angle of the inlet velocity in the device. For fully developed flows, the components of the WSS vectors at the device centroid are given in terms of the dynamic viscosity, *μ*, the channel height, ℎ, and the mean flow velocity components in the X and Y directions, *V_X_*_|*C*_and *V_Y_*_|*C*_ as

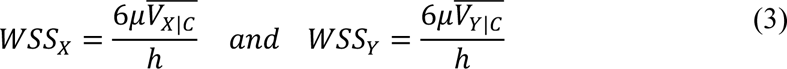

These mean flow velocities at the device centroid are calculated using the inlet velocities from the continuity equation as

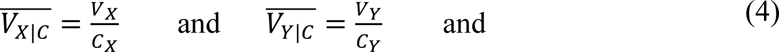

*C_X_* and *C_Y_* are channel-specific coefficients, which are essential to compute the increase in area at the centroidal region as compared to the areas at the inlet ports connected to the syringe pumps. For the device geometry fabricated in this study, *C_X_* = 15 and *C_Y_* = 14. Other geometries will result in different values of *C_X_* and *C_Y_* in these equations. *V_X_* and *V_Y_* are the X and Y components of the resultant velocities that are obtained by vectorially adding the inlet velocities, *V*_1_and *V*_2_.

The ratio of WSS components at the centroid is written in terms of the phase angle of the inlet velocity in the device, *η*, by substituting Equations 2 and 4 in 3 as

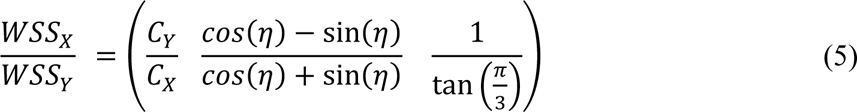

The magnitude of velocity, *V_0_*, obtained by substituting 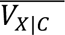 (Equation 4) in the expression for *WSS_X_* (Equation 3), is given by

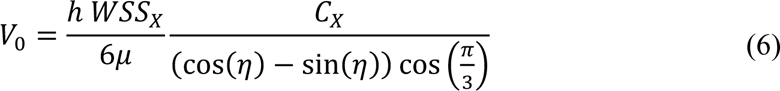

These equations are used to calculate the velocity inputs to obtain any targeted rosette shape in the device with known values of *WSS_X_* and *WSS_Y_* and known area increment coefficients. *V_0_* and *ƞ* may be calculated at each time instant, t, using Equations 5 and 6. We use these expressions in Equation 2 to calculate the inlet velocitiesat the syringe pumps using the ‘sum of sine fit’ and obtain variations in *V*_1_ and *V*_2_ with time. For the circular rosette (Figure 1d), with a WSS magnitude of 1 Pa, the predicted *V*_1_ = *V*_2_(*t*)and *V*_2_ = *V*_2_(*t*) are given using Equations 1. WSS magnitude of 1 Parepresents the physiological shear stresses experienced by endothelial cells due to blood flows.^15^

We used a commercial computational fluid dynamics solver (ANSYS-Fluent, 2020 R1) to simulate unsteady, viscous, and laminar flows in the channel using velocity inputs to the pump given by the semi-analytical model in Equation 1 for a circular rosette. The CFD model uses pressure-velocity coupling with laminar flow conditions, and a first-order implicit method to discretize the time-step at 1 ms **(Figure S1).** Spatial discretization of the convection and gradient terms was performed using the 2^nd^ order upwind and least square cell-based methods, respectively. The residual criteria for three velocity components and mass conservation were set to 1E-04. A mesh resolution of 10 cells along the height direction was selected from grid independence studies **(Figure S2 and Table S1).** 21,12,370 elements were included in the device geometry. Fluid density, *ρ* = 1055 kg/m^3^, and dynamic viscosity, *μ* = 0.0049 kg/ms, were used in these computations. Wall boundary conditions were implemented on all microchannel surfaces that connect both inlets to the outlet of the device. A velocity inlet boundary condition was used at both inlets, and a pressure outlet boundary condition was used for the reservoir. Post-processing was performed using Paraview software.

WSS variations in the device were obtained from CFD simulations, and the results were compared with the semi-analytical model to validate this method. We calculated the Anisotropy Ratios (AR) at each wall node in the device geometry to generate contour plots from the CFD analysis for each of the different targeted shear rosettes. The contour plots allowed us to visualize and understand the distribution of AR values across the device. This validation is centralin studying the effects of complex flow conditions on endothelial morphology, signaling, and transduction responses.

### 4.2 Fabrication and validation of microfluidic devices to generate controlled flows

We used a replica molding technique to prepare microfluidic devices in this study using chemically etched sheets of stainless steel (100 µm). In this method, we attached an etched sheet to a glass slide treated with trichloro silane (1H,1H,2H,2H-perfluorooctyl; Sigma Aldrich, India) in a desiccator overnight. Polydimethylsiloxane (PDMS; Sylgard®184, Dow Corning, USA) was mixed in a stoichiometric ratio of 10:1 based on the manufacturer’s instructions, poured over the prepared channel geometries, degassed to remove air bubbles, cured at 80°C for 2 hours, and peeled carefully to obtain the microfluidic channels. The channel height was 100 μm, and the width of the cross-sections in the three arms was 200 μm. We used a similar procedure to fabricate straight microfluidic channels using laser-cut acrylic tape with dimensions of 10 mm length X 1 mm width X 100 µm height for experiments using unidirectional laminar (Uni-L) and oscillatory (Uni-O) flows. Inlet and outlet holes were punched using a 0.5 mm biopsy punch, and the devices were cleaned and sterilized with 100% ethanol for cell culture experiments. Molded PDMS parts were clamped to coverslips using a 3D-printed hollow frame to reduce lateral movement, and cells were visualized under an inverted microscope during the experiment. The frame was pressed down by four screws to apply uniform pressure on the PDMS channel and coverslip, this method was useful to prevent leaks during the flow experiments.^29^ Two synchronized syringe pumps allow liquid at desired flow rates,derived for the specific rosette shape as described earlier, into the device, and produce oscillatory or steady flows within the central region. The third arm of the microfluidic channel in the device was connected to a reservoir. Absence of moving parts makes the device robust in utility, durability, and replicability. The frame was dismantled at the end of the flow experiment, and the cells were fixed on the coverslip for immunofluorescence studies.

We performed flow visualization experiments using 0.1% v/v green fluorescent 2 µm sized polystyrene beads in MilliQ water astracers (Movie S3). Beads were injected into the channel area using syringe pumps and the flow was visualized after a settling time of 15 minutes. Images were captured on the Leica SP5 epifluorescence system at 25 frames per second **(Movie S3)**. The streak images indicate streamlines that closely match with streamlines obtained using computation.

### 4.3 Cell Culture and Seeding

Human aortic endothelial cells (HAECs) were purchased from LONZA (Lonza cat. no. CC-2535) and used at early passages of 2 to 5 in all experiments. Cells were cultured in EGM™-2 Endothelial Cell Growth Medium-2 BulletKit (Lonza cat. no. CC-3162) in sterile conditions within a 5% CO_2_ and 37°C incubator. Glass coverslips (22 mm) were cleaned using ultrasonication, activated using oxygen plasma for 2 minutes, and coated with 40µg/ml of rat tail collagen-1 (A1048301, Thermo) at 37°C for 1 hour in a humid chamber. Prepared coverslips were washed three times with phosphate-buffered saline (PBS, Thermo) and seeded with HAECs at 1 × 10^6^ cell/mL density to produce a ∼90% confluent EC monolayer in 48 hours.

The PDMS channel was placed on the coverslip with the HAEC monolayer, and the channel was press-fiton the frame as described earlier. This method differs from other studies that take a significantly long time to prepare channels with confluent endothelial cells for flow studies. These earlier procedures involve flowing a small volume of media into microfluidic channels cultured with endothelial cells, waiting for cell attachment and monolayer confluency within the chamber, and then performing the flow studies. We used programmable syringe pumps (NE-1000, Syringepump.com) that were synchronized using an RS-232 dual pump communications cable. In this method, one pump serves as the Master control pump and the other as the secondary slave. Inlet flow rates from Equation 1 were calculated for various time points over the entire cycle and were input to the two pumps. Experiments on unidirectional laminar (Uni-L) and oscillatory (Uni-O) experiments were performed for 24 hours using a single pump with a duty cycle similar to the Bi-O studies.

### 4.4 Immunofluorescence studies

Cells were washed with PBS buffer (Fisher Scientific), fixed with 4% paraformaldehyde for 15 minutes, permeabilized with 0.1% Triton X-100 for 3 minutes, and blocked in 5% BSA + 5%FBS for 1 hour at room temperature for immunofluorescence studies. Samples were incubated for 1 hour at room temperature in primary antibody for VE-Cadherin (1:500, 2158, CST), Lamin A/C (1:500, MA3-1000, Thermo), and secondary Alexa fluor 488 (1:600, Molecular Probes, Invitrogen) antibody and Rhodamine-phalloidin (1:200, Invitrogen) for 1 hour at room temperature, and rinsed twice using PBS. Samples were treated with DAPI (1:500, Invitrogen) for 2 minutes, rinsed twice with PBS to label nuclei, and mounted on coverslips with Goldfade (Invitrogen). A Leica SP5 FALCON confocal microscope was used to image the samples. Lamin A/C was imaged at the nucleus midsection and the entire cell stack was acquired for Actin, VE-Cadherin, and DAPI to report differences in fluorescence values. Cell areas and aspect ratios were quantified using the Tissue analyzer (Fiji by ImageJ).

### 4.5 RNA Isolation and RT-QPCR

Cell monolayers were washed with PBS buffer (Fisher Scientific) and treated with 200µl of TRIzol® reagent (ThermoScientific) per coverslip in a PDMS well of an internal diameter of 1.5mm placed on the device centroid to isolate the cells subjected to 0.6 ≤ *AR* ≤ 1.0 for analysis in the Bi-O condition. Cells subjected to Uni-L and Uni-O groups were isolated and collected from the channels. RNase Mini Kit (Qiagen) was used per the manufacturer’s instructions. 2 µg of RNA was used for cDNA synthesis using a high-capacity cDNA reverse transcription kit (Applied Biosystems). PCR products were checked using Quantitative real-time PCR (qRT-PCR) using a Power up SYBR Green master mix (ThermoScientific) with 10ng of cDNA as the template. Gene expressions were normalized to GAPDH. Fold change was calculated using the 2^-ΔΔct^ method and normalized again to (Control) No-flow groups. Gene expressions were analyzed across three biological replicates in this study.

### 4.6 Statistical analysis

Experiments were performed in three independent trials. Data are reported as mean ± SD. Results from the different groups were compared using a one-way analysis of variance (ANOVA) with Bonferroni comparison to test for individual differences. p< 0.05 is indicated with (*).

## 5.0 Acknowledgements

We thank the Bioimaging Facility, Division of Biological Sciences, IISc, for the use of the Lecia SP8 FALCON confocal microscope. We also acknowledge the early contributions of Mr. Vamsi Krishna in this work. JHA and NG are grateful to DBT (BT/PR48494/MED/32/866/2023) for project support.

## 6.0 Author contributions

Conceptualization: JHA, NG; Methodology: NP, SM, JHA, NG; Software & Modeling: SM; Validation: NP, SM; Formal Analysis: NP, SM, JHA, NG; Investigation: NP, SM; Data Curation: NG; Writing - Original Draft Preparation: NP, SM, NG; Writing - Review & Editing: NP, SM, JHA, NG; Visualisation: NP, SM, NG; Supervision: JHA, NG; Funding Acquisition: NG.

---

**Table S1:** Mesh size independence study in the CFD study used the following parameters.

|  |  |  |  |
| --- | --- | --- | --- |
| **No. of cells along the height** | 05 | 07 | 10 |
| **Minimal cell size [$\mu\text{m}$]** | 20 | 15 | 10 |
| **\|WSS\| [Pa]** | 1.91 | 1.99 | 2.03 |

**Table S2:**
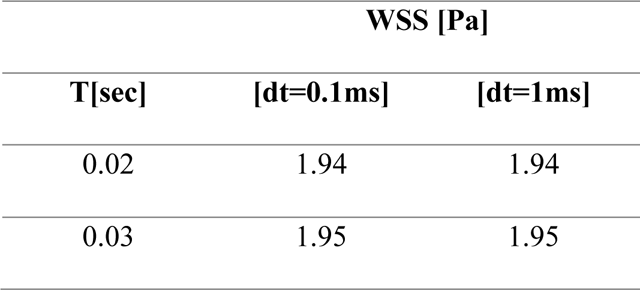
Time step selection for CFD calculations used the following parameters.

**Figure S1.**
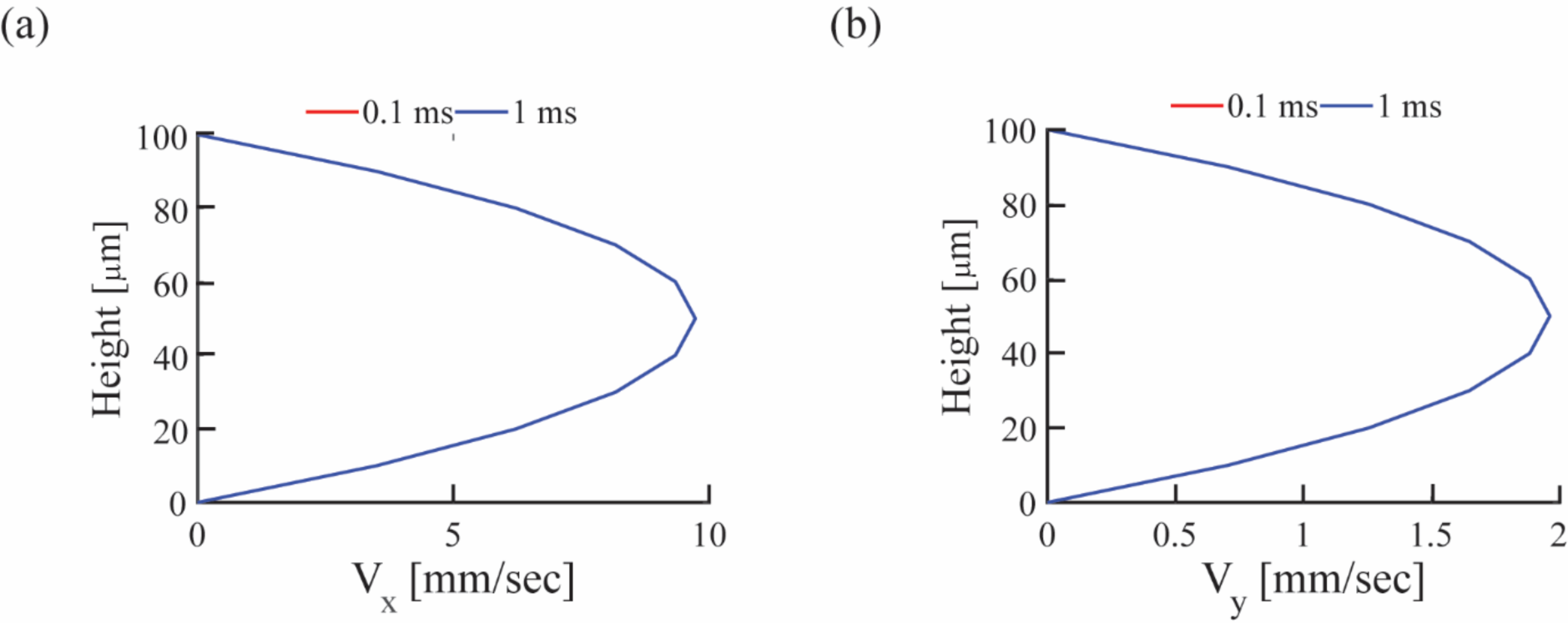
**(a):**X and **(b)** Y components of the velocity profiles were extracted at the device centroid at 30 ms for the circular shear rosette. Because the velocity profiles for 1 ms and 0.1 ms are same, we selected a higher time step of1 ms for faster final computations. WSS magnitude at the centroid of the channel was maintained to be a constant for these calculations (**Table S2**).

**Figure S2:**
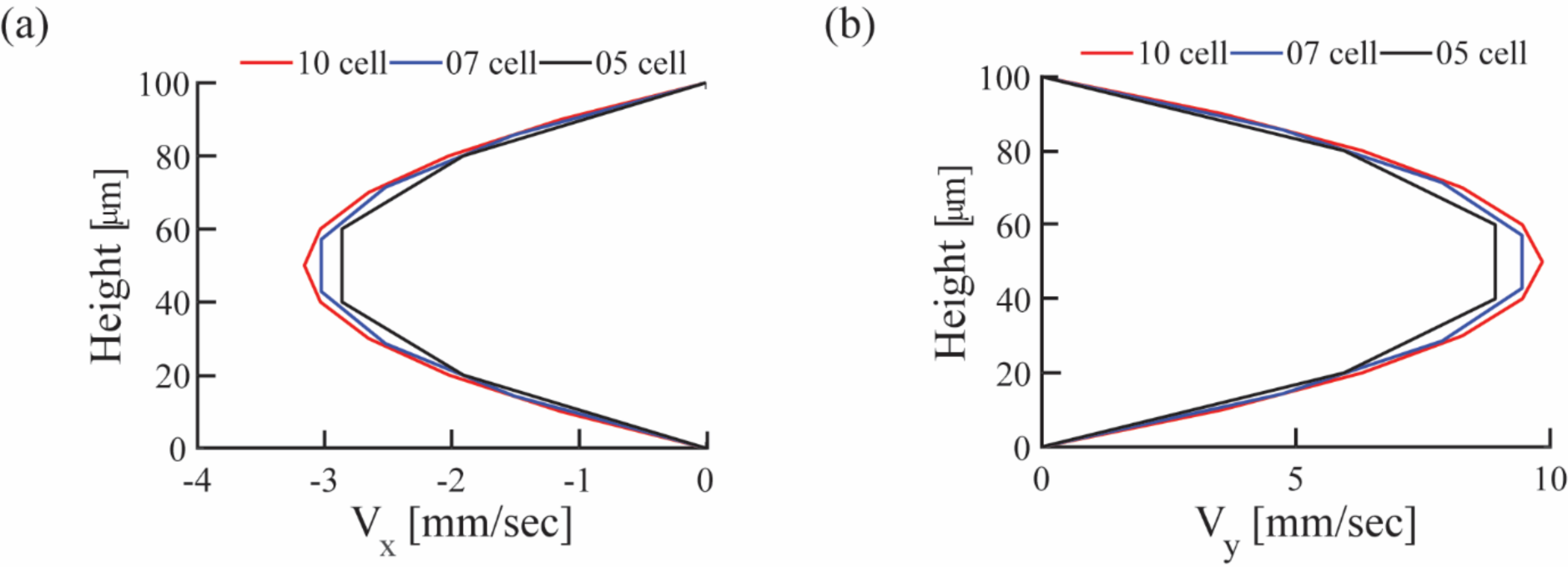
To assess mesh independence, we used 5, 7, and 10 cells across the height of the channel, and plotted **(a)** X componentof the velocity, and **(b)** Y component of the velocity for a circular shear rosette. Velocities were extracted at the centroid of the channel at 300 ms. A mesh resolution of 10 cells showed comparatively smooth velocity profiles, and was used for the final computations. The magnitude of WSS at the centroid of the channel is near constant for simulations corresponding to 7 and 10 cell meshes (**Table S1**).

**Figure S3.**
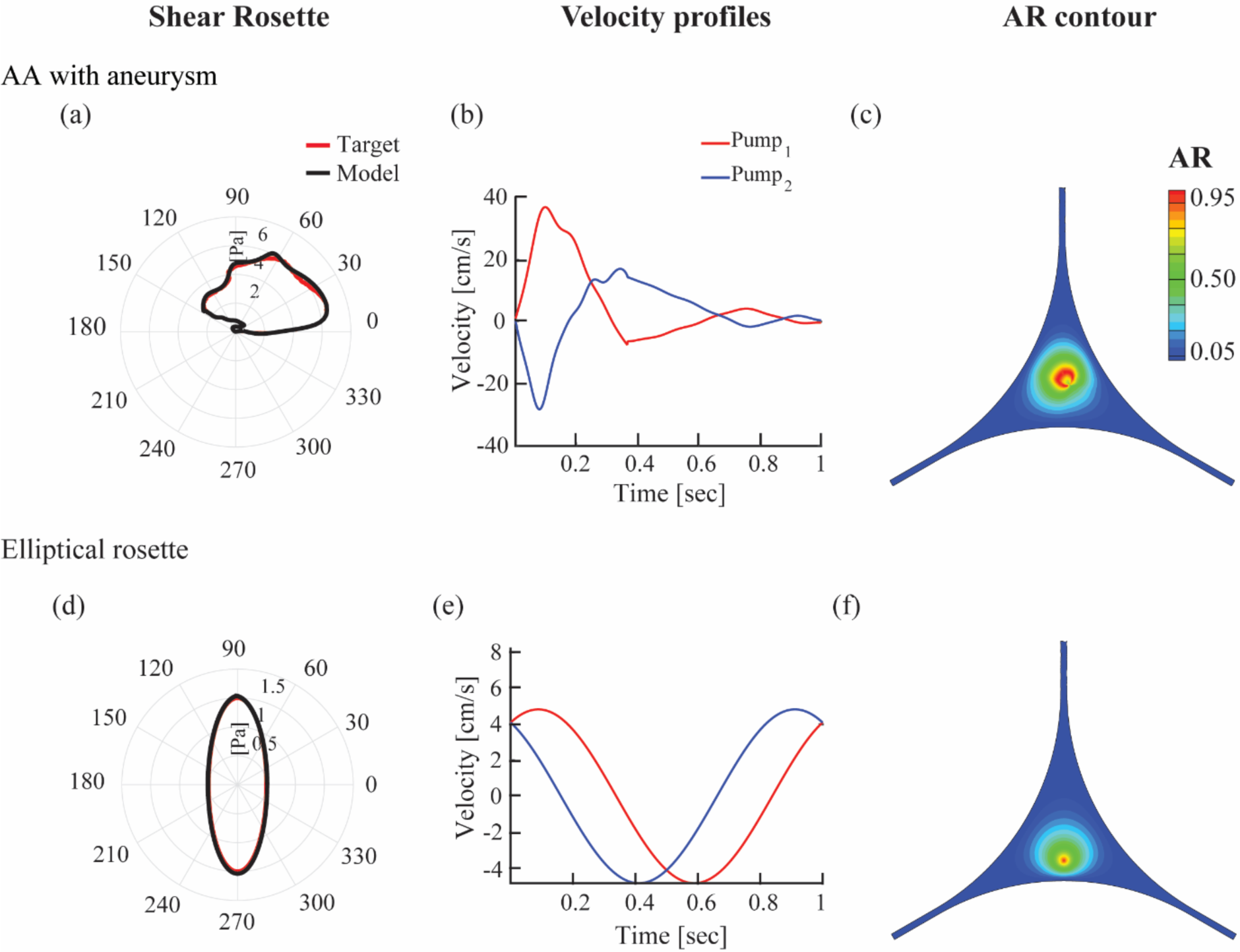
**(a)** The semi-analytical model in our study may be used to generate any rosette shape in disturbed flow regions. Results showa close match between a target rosette (red) for a point in an aortic aneurysmal (AA) regionanda computationally obtained rosette (black) at the centroid of the device using the semi-analytical model.^15,29^ **(b)** The corresponding velocity profiles from the semi-analytical model served as inputs to syringe pumps in the device. **(c)** Contour plots show AR distributions for the AA rosette. **(d)** We note a good match between results from a target elliptical rosette (red) of magnitude 1 Pa with calculated rosette (black) obtained using the semi-analytical model. **(e)** Corresponding velocity inputs from the model to generate the elliptical rosette serve as inputs to the two syringe pumps. **(f)** The AR distribution is shown in the device for the elliptical rosette.

**Movie S1.**
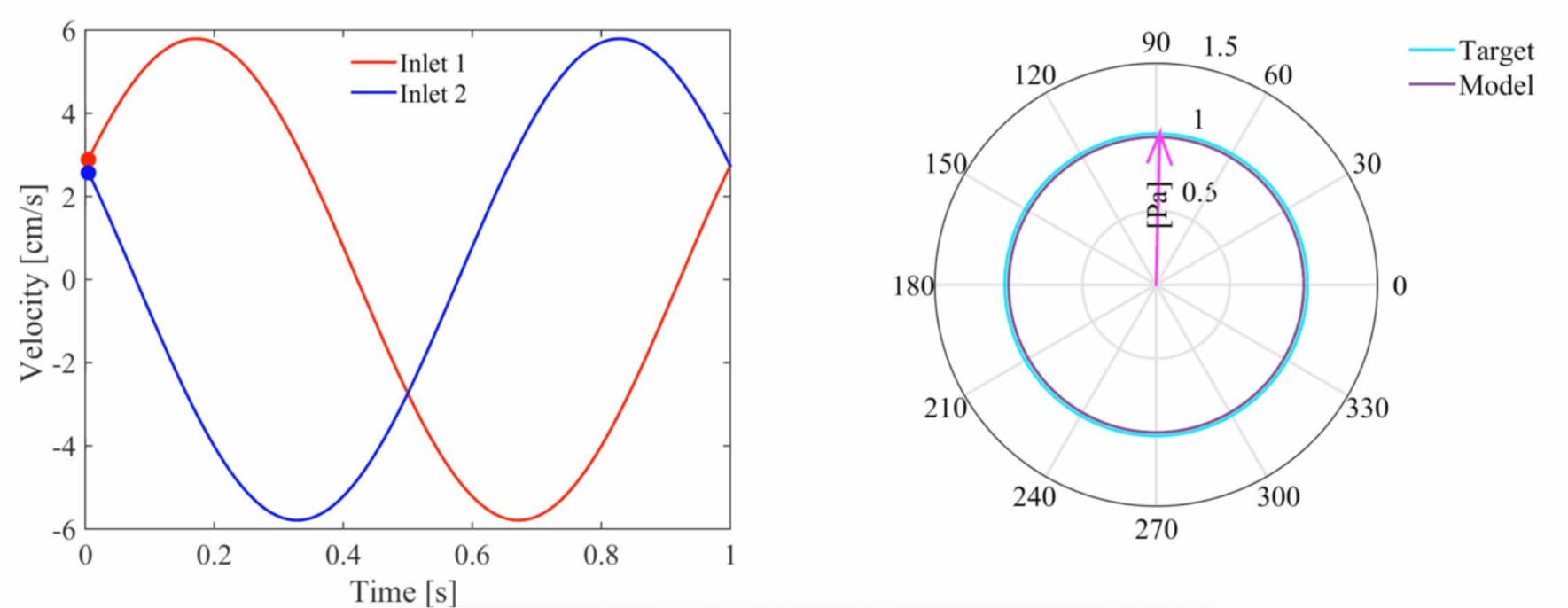
**Title:** Inlet velocity variations with time are shown for a circular rosette shape. **Description:** Velocity inputs to the two pumps connected to the inlet ports of the endothelium-on-chip device were obtained using the semi-analytical model for a circular shear stress rosette at the device centroid.

**Movie S2.**
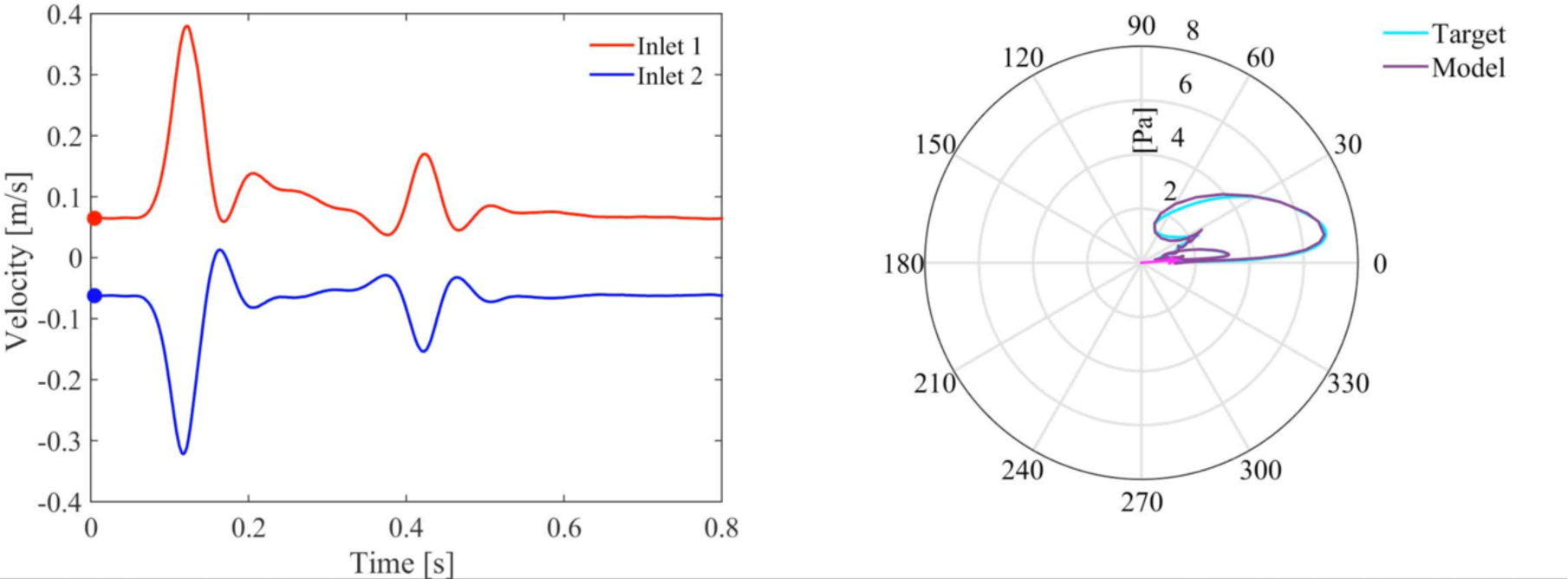
**Title:** Inlet velocity variations are shown for a rosette in adiseased carotid artery. **Description:** The velocity inputs and corresponding WSS variations in the endothelium-on-chip device are illustrated for a shear rosette pattern corresponding to an internal carotid artery (ICA) with an aneurysm^12^ reported in an earlier study.

**Movie S3.**
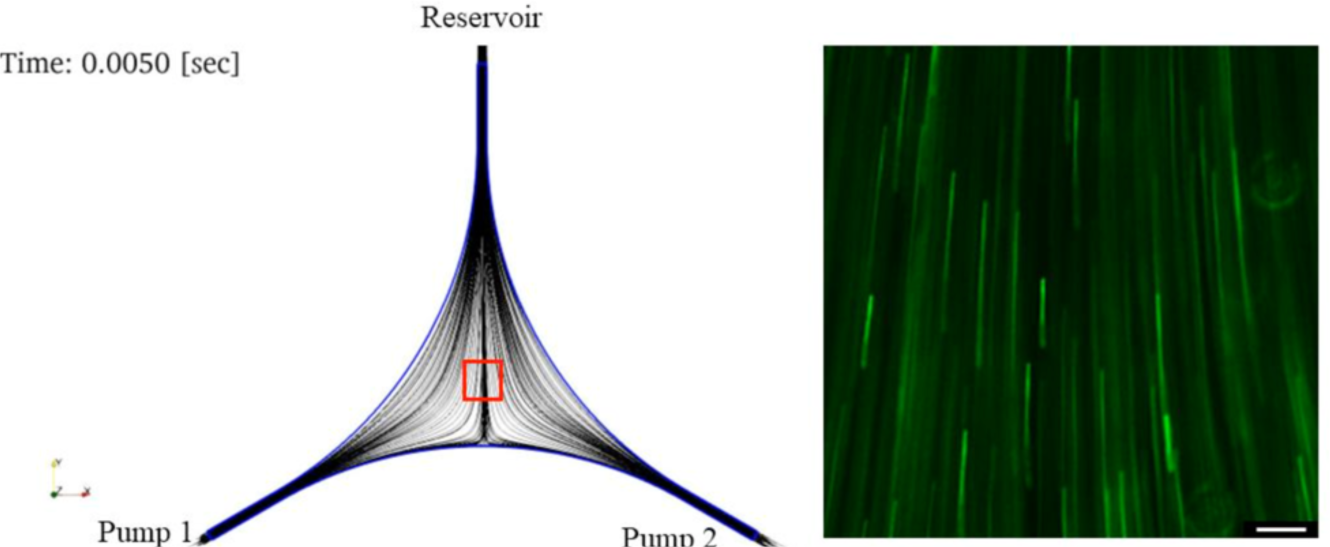
**Title:** Comparisons between streamlines from CFD studies with flow visualization studies. **Description:** Computationally obtained streamlines, depicting fluid flows in the endothelium-on-chip device, match closely with streak images obtained from experimental flow visualization, performed with 2 μm tracer beads suspended in the fluid, at a region in the device identified with a red box.

