## supplementary table for "Endothelial mechanobiology under controlled disturbed flows"

**Table T1**: Mesh independence study: size

| No. of cells along the height | 05 | 07 | 10 |
| --- | --- | --- | --- |
| **Minimum cell size [µm]** | 20 | 15 | 10 |
| **\|WSS\| [Pa]** | 1.91 | 1.99 | 2.03 |

**Table T2**: Time step selection

| | WSS [Pa] | |
| --- | --- | --- |
| **T[Sec]** | **[dt=0.1mSec]** | **[dt=1mSec]** |
| 0.02 | 1.94 | 1.94 |
| 0.03 | 1.95 | 1.95 |
